# *Phytophthora cinnamomi* populations collected from avocado in the United States exhibit high adaptive capacity to climate and disease control methods

**DOI:** 10.64898/2026.04.28.721487

**Authors:** Benjamin K. Hoyt, Savannah Salas, Jonathan H. Crane, Monica Navia-Urrutia, Romina Gazis, Liliana M. Cano, Achyut Adhikari, Miaoying Tian, John Jifon, Ricardo Goenaga, Luz M. Serrato-Diaz, James E. Adaskaveg, Patricia Manosalva

## Abstract

*Phytophthora cinnamomi*, the causal agent of Phytophthora root rot (PRR), poses a persistent threat to the United States avocado industry, the top domestic producer and consumer. Avocado growers are facing clonal A2 *P. cinnamomi* populations challenging their current PRR control methods. In this study, we characterized 125 isolates collected from orchards in California, Florida, Hawaii, Texas, and Puerto Rico for radial growth per day, optimal growth temperature, *in vitro* fungicide sensitivity, and virulence on D’Anjou pear fruit and UC2001 avocado seedlings. Across all isolates, optimal growth occurred most frequently at a range from 22 to 25°C; however, a subset of isolates from Hawaii, Florida, and California exhibited higher optimal growth temperatures (28°C and 30°C) suggesting thermal adaptation in warmer regions. Potassium phosphite EC_50_ values spanned from 4.61 to 763.13 µg/ml, with significantly higher insensitivity in isolates from California and Florida, reflecting the continued overuse of this fungicide in these major production states. In contrast, baseline sensitivities to ethaboxam, mandipropamid, mefenoxam, fluopicolide, and oxathiapiprolin were uniformly high, with narrow, unimodal EC_50_ distributions across states. Finally, a wide range of virulence among isolates was detected using avocado seedlings and D’Anjou pear fruits with isolates from California and Puerto Rico being the most virulent. Together, this data documents extensive phenotypic diversity within clonal A2 *P. cinnamomi* populations including heat-adapted and phosphite-insensitive lineages, establishes multi-state fungicide sensitivity baselines, and underscores the need for continued surveillance, integrated fungicide stewardship (especially phosphonates), and rootstock screening against phenotypically diverse populations to sustain avocado PRR management and ensure the United States’ avocado industry sustainability and profitability.

## 1 Introduction

Avocado (*Persea americana*, Mill) is a major fruit tree crop produced worldwide. The avocado fruit is a globally popular, nutrient-dense superfood with numerous health benefits (Dreher and Davenport, 2013). In 2023, the global avocado production reached 10 million metric tons with Mexico being the top producer and exporter (FAO, 2025). *Phytophthora cinnamomi*, the causal agent of avocado Phytophthora root rot (PRR), is a highly invasive soilborne oomycete pathogen recognized as a major threat for agriculture as well as global biodiversity (Burgess et al., 2017). Its exceptionally broad host range, infecting more than 5,000 plant species, has led to its inclusion among the world’s top 100 invasive alien species (Lowe et al., 2000). Because PRR is one of the most destructive diseases affecting the sustainability of the avocado industry, it has been the focus of extensive research in major producing regions including the U.S.A. (Belisle et al., 2019a; Crandall, 1948; Shands et al., 2025; Zentmyer, 1976), Mexico (Mondragón-Flores et al., 2022; Ochoa-Fuentes et al., 2007; Zentmyer, 1951), and South Africa (Engelbrectht et al., 2022; Köhne, 1998; Wager, 1942). As global avocado cultivation has expanded to every continent except Antarctica, reports of PRR outbreaks and studies on *P. cinnamomi* have increased markedly over the last decade (Kavroulakis et al., 2024; Kurbetli et al., 2020; Madhu et al., 2025; Ramírez-Gil et al., 2021).

The U.S.A. is one of the world’s largest avocado consumers and its top importer (FAO, 2025; Hammami et al., 2024). Domestically, California (CA) remains the leading producer, followed by Florida and Hawaii (Hammami et al., 2024). PRR causes substantial economic losses through reduced yields, diminished fruit quality, and the need for costly disease management practices (Coffey, 1987; Ramírez-Gil et al., 2017). Disease severity is further intensified by environmental conditions common in U.S.A. producing states, including waterlogged soils, increased hurricane frequency and flooding in Florida and Puerto Rico, heavy rainfall in Hawaii, and widespread pathogen presence in Texas orchards; all factors that constrain the U.S.A. industry expansion (Bender et al., 2012; Hawaii Department of Agriculture, 2024; Kunta et al., 2007; Ploetz and Schaffer, 1989; RoyChowdhury et al., 2016).

*Phytophthora cinnamomi* is a heterothallic hemibiotrophic pathogen requiring two mating types (A1 and A2) for sexual reproduction; however, only clonal A2 populations have been associated with avocado PRR worldwide (Calle-Hanao et al., 2020; Dobrowolski et al., 2003; Engelbrecht et al., 2022; Hüberli et al., 2001; López-Herrera and Pérez-Jiménez, 1995; Shands et al., 2025). The pathogen reproduces primarily through its asexual cycle: sporangia release motile zoospores that move through moist or waterlogged toward avocado roots by chemotaxis (Hardham and Blackman, 2018). Under unfavorable conditions, thick-walled chlamydospores allow this pathogen long-term survival (Jung et al., 2013). In orchards, *P. cinnamomi* spreads through soil, water, and infested plant material, while human-mediated movement of contaminated soil, nursery stock, or equipment enables long-distance dispersal (Dreistadt, 2007). Infections of feeder roots, and occasionally trunk cankers, lead to canopy decline, reduced yields, and often tree mortality (Hardham and Blackman, 2018; Ploetz and Parrado, 1987). Entirely newly planted orchards may fail in the absence of effective disease management (Faber et al., 2012). Current PRR management includes chemical treatments (e.g., phosphonates, mefenoxam, and oxathiapiprolin) (Adaskaveg et al., 2024), tolerant or resistant rootstocks (Bender et al., 2012; Crane et al., 2025), accurate diagnostics (Wilkins et al., 2025), and cultural practices such as mulching and irrigation management (Ramírez-Gil and Morales-Osorio, 2020). However, reports of *P. cinnamomi* isolates exhibiting reduced sensitivity to potassium phosphite and increased virulence on widely used avocado rootstocks such as Dusa^®^ and Duke7 have risen in the past decade (Andronis et al., 2024; Belisle et al., 2019a; Belisle et al., 2019b; Dobrowolski et al., 2008; Hoyt et al., 2026; Hunter et al., 2023; Ma and McLeod, 2014; Sánchez-González et al., 2019; Shands et al., 2025). These findings align with the substantial phenotypic diversity documented in *P. cinnamomi* populations from avocado orchards in Mexico (Mondragón-Flores et al., 2022; Ochoa-Fuentes et al., 2007), Colombia (Calle-Henao et al., 2020; Palacios-Joya et al., 2024), New Zealand (Hunter et al., 2023), Australia (Wilkinson et al., 2001), and South Africa (Ma and McLeod, 2014).

In California, our previous work identified two clonal A2 lineages associated with PRR, both displaying broad phenotypic variation in radial growth rate, optimal growth temperature, fungicide sensitivity, and virulence (Belisle et al., 2019a; Belisle et al., 2019b; Shands et al., 2025). Pathogen isolates collected from southern growing regions exhibited less sensitivity to higher growth temperatures and potassium phosphite but at the same time exhibited more virulence when infecting commonly used avocado rootstocks (Belisle et al., 2019a; Belisle et al., 2019b; Shands et al., 2025). Moreover, we found that *P. cinnamomi* isolates corresponding to one of the A2 clonal lineages detected only in Southern California growing regions (Pagliaccia et al., 2013) were genetically similar to *P. cinnamomi* populations collected from avocado orchards in Mexico (Mexican origin) (Ochoa-Fuentes et al., 2015) suggesting migration between the two regions (Shands et al., 2025). Despite the importance of avocado PRR to U.S.A avocado production, comprehensive genotypic and phenotypic characterization of *P. cinnamomi* in other producing states remains limited. To address this gap, we expanded the phenotypic characterization of *P. cinnamomi* populations from avocado orchards in Florida, Hawaii, Texas, and Puerto Rico, examining *in vitro* growth rates, optimal growth temperatures, *in vitro* fungicide sensitivity, and virulence. We also evaluated phenotypic shifts in the current California populations. Continued monitoring of domestic pathogen populations is essential to guide the development, evaluation, and long-term durability of PRR management strategies, ultimately supporting the sustainability, competitivity, and profitability of the U.S.A. avocado industry.

## 2 Materials and methods

### 2.1 Collection of *Phytophthora cinnamomi* isolates

*Phytophthora cinnamomi* isolates from California, Florida, Hawaii, Texas, and Puerto Rico (2020-2022) were recovered from symptomatic avocado trees and soil samples using multiple isolation methods. Root and soil plating were performed as described by Belisle et al. (2019a), and soil baiting following the protocols of Hao et al. (2018) using pear fruits or avocado leaves. Isolates were identified morphologically, and single-zoospore cultures were generated following Lonsdale et al. (1988). Molecular confirmation was conducted by Sanger sequencing of the internal transcribed spacer (ITS) region and a *P. cinnamomi*-specific TaqMan quantitative PCR assay targeting the mitochondrial *atp9*-*nad9* locus (Bilodeau et al., 2014; Belisle et al. 2019a; Miles et al., 2017). In addition, four isolates collected from diseased walnut trees in California were provided by Dr. Gregory T. Browne (University of California, Davis) (Browne et al., 2015). All isolates were stored as colonized 10% clarified V8 (V8C) agar plugs in water (Ko, 2003) until use.

### 2.2 *In vitro* radial growth rate and optimal growth temperature

Isolates were assessed for *in vitro* radial growth rate per day (RGPD) and optimal growth temperature on 10% V8C agar at 22°C, 25°C, and 28°C following Shands et al. (2025). For isolates that continued to increase in growth at 28°C, additional measurements were obtained at 30°C and 32°C. The previously characterized isolate, Pc2113, was included as an internal control (Shands et al., 2025).

Experiments were performed in triplicate and repeated twice.

### 2.3 *In vitro* sensitivity to Oomycota fungicides

Sensitivity to potassium phosphite (Prophyt, 34.3% phosphorous acid [FRAC code P07]; Helena Chemical Co., Collierville, TN) was conducted using the agar dilution method described in Adaskaveg et al. (2015) with potassium phosphite concentrations of 0 (control), 5, 25, 100, 150, 300, 600, or 1000 μg/ml. EC_50_ values (inhibition of 50% mycelial growth) were calculated as described in Shands et al. (2025).This experiment was performed in triplicate and repeated twice. *In vitro* sensitivity to mefenoxam (Ridomil Gold^®^ SL [FRAC code 4]; Syngenta Crop Protection, Greensboro, NC), oxathiapiprolin (Orondis^®^ [FRAC code 49]; Syngenta Crop Protection, Greensboro, NC), mandipropamid (Revus [FRAC code 40]; Syngenta Crop Protection, Greensboro, NC), fluopicolide (Presidio [FRAC code 43]; Valent U.S.A., Walnut Creek, CA), and ethaboxam (Elumin [FRAC code 22]; Valent U.S.A., Walnut Creek, CA) was done using the spiral gradient dilution method (SGD) described by Belisle et al. (2019b). Aqueous stock solutions of mefenoxam (50 µg/ml), oxathiapiprolin (1 µg/ml), mandipropamid (10 µg/ml), fluopicolide (100 µg/ml), and ethaboxam (50 µg/ml) were applied to 15-cm 10% V8C agar plates using a spiral plater (Eddy Jet 2W, Neutec Group, Inc., Farmingdale, NY) in exponential deposition mode. Sterile Milli-Q water (Millipore Sigma, Burlington, MA, USA) was used for control plates. Two mycelium-covered cellophane strips per isolate were placed opposite each other on each chemical-or water-amended plates and incubated at 22°C for 2 days. The point of 50% growth inhibition was recorded and EC_50_ values were calculated using the Spiral Gradient Endpoint (SGE) software (Spiral Biotech). Pc2113 was included as an internal control (Belisle et al., 2019b; Shands et al., 2025). Experiments were repeated twice.

### 2.4 Virulence assays

Virulence was assessed on D’Anjou pear fruit as described by Shands et al. (2025). Six fruits per isolate were inoculated with 20 µl of a 1×10^4^ zoospore/ml suspension. Lesion area was recorded at 3-, 4-, and 5-days post inoculation (DPI) to calculate area under the disease progression curve (AUDPC). Experiments were repeated twice and Pc2113 served as an internal control (Shands et al., 2025). Virulence on avocado was evaluated using UC2001 seedlings (Menge et al., 1999) following Shands et al. (2025) with modifications. Avocado seedlings were grown for 12-14 weeks at temperatures and humidity conditions ranging from 22°C to 30°C and 35% to 72%, respectively. For each isolate, five seedlings were inoculated with 3.4 g of *P. cinnamomi*-colonized millet, and the percentage of diseased roots was assessed at 5 weeks post-inoculation (WPI). Pc2113 and Pc2109 were included as internal controls (Shands et al., 2025). Seedlings treated with uninoculated millet served as negative controls.

### 2.5 Statistical Analysis

Data normality was assessed for using the Shapiro-Wilk test. Except for seedling virulence assays, all datasets were analyzed using generalized linear mixed models (GLMMs) with a gamma distribution, log link function, and experimental repeats as random effects. Seedling virulence data were analyzed using a generalized linear model (GLM). Least squares means were calculated using LSMEANS with a TUKEY adjustment. Analyses were performed in R v.4.3.2 (R Core Team 2023) using the lme4 (Bates et al., 2015) and lsmeans (Lenth, 2016) packages. Results were considered significant at *P* ≤ 0.05.

## 3 Results

### 3.1 *P. cinnamomi* populations from California, Florida, and Hawaii exhibit adaptation to higher temperatures

A total of 125 *P. cinnamomi* isolates were examined in this study, including four isolates from walnut in California. Avocado isolates originated from California (n = 34), Florida (n = 25), Hawaii (n = 12), Texas (n = 25), and Puerto Rico (n = 25) (Table 1). The RGPD values for all isolates at each growth temperature tested are summarized in Supplementary Table S1. Across all isolates, optimal growth temperatures occurred at 22°C (52.8%) and 25°C (31.2%), while only 13% growing optimally at 28. Previously, we reported that the *P. cinnamomi* population (1994-2019) from southern growing areas in California (San Diego and Riverside Counties [CA-South]) displayed greater virulence and enhanced growth at warm temperatures compared to isolates from northern growing areas (Ventura and Santa Barbara [CA-North]) (Shands et al., 2025). Here, we observed similar patterns in the current population (2020-2022). CA-North isolates grew significantly faster than CA-South isolates at 22°C and 25°C, but not at 28°C. Consistent with this, 53% and 47% of CA-North isolates exhibited optimal growth temperatures of 22°C and 25°C, respectively, whereas 41% of CA-South isolates grew optimally at 28°C and 29% at 22°C and 25°C. The average RGPD of CA-North isolates decreased significantly at 28°C compared with 22°C and 25°C, while CA-South isolates maintained similar growth across temperatures (Supplementary Table S2 and Supplementary Figure S1).

**Table 1.** Sample information of all *Phytophthora cinnamomi* isolates used in this study.

| Number of Isolates | State | County | Host | Year Collected (range) |
| --- | --- | --- | --- | --- |
| 7 | California | Riverside | Avocado | 2020-2022 |
| 10 | California | San Diego | Avocado | 2020-2022 |
| 12 | California | Santa Barbara | Avocado | 2020-2022 |
| 5 | California | Ventura | Avocado | 2020-2022 |
| 9 | Florida | Miami-Dade | Avocado | 2020-2021 |
| 11 | Florida | Lee | Avocado | 2022 |
| 2 | Florida | St. Lucie | Avocado | 2022 |
| 3 | Florida | Polk | Avocado | 2022 |
| 2 | Hawaii | Honolulu | Avocado | 2021 |
| 7 | Hawaii | Hawaii | Avocado | 2021-2022 |
| 3 | Hawaii | Kauai | Avocado | 2022 |
| 7 | Texas | Cameron | Avocado | 2021 |
| 18 | Texas | Hidalgo | Avocado | 2022 |
| 9 | Puerto Rico | Adjuntas | Avocado | 2021-2022 |
| 16 | Puerto Rico | Isabela | Avocado | 2022 |
| 1 | California | Colusa | Walnut | 2010 |
| 1 | California | Merced | Walnut | 1996 |
| 2 | California | San Joaquin | Walnut | 2009 |

Significant isolate-level differences in RGPD were detected at all temperatures. As expected, control isolate Pc2113 grew optimally at 22°C. The slowest-growing isolates at 22°C and 25°C were from Florida (AVR-001 and AVR-008), and at 28°C from Hawaii (HI-8-SZ1). All fastest-growing isolates originated from California (Figure 1; Supplementary Table S1). Significant differences in average RGPD among states were detected at 25°C and 28°C, with isolates from Puerto Rico exhibiting the lowest values compared with those from California, Florida, and Hawaii (Figure 1; Supplementary Table S3). Most isolates from Texas (76%) and Puerto Rico (80%) had an optimal growth at 22°C, and none grew optimally above 25°C. Twenty isolates from California, Florida, and Hawaii did not exhibit reduced growth at 28°C; prompting further evaluation at 30°C and 32°C. Of these 20 isolates, 16 grew best at 28°C. Seven of eight isolates from California with optimal growth at 28°C originated from CA-South (Riverside and San Diego) while four of nine Florida isolates grew optimally at 30°C (Table 2 and Figure 2). These results highlight the adaptive capacity of *P. cinnamomi* populations to higher temperatures in Southern California growing regions, Florida, and Hawaii.

**Table 2.** Optimal growth temperature of select *Phytophthora cinnamomi* isolates exhibiting increasing *in vitro* radial growth per day (RGPD) beyond 28°C on 10% V8C (clarified V8) agar. Preferred average optimal RGPD values, standard deviation, and temperature are displayed for each isolate.

| Isolate ID | State | Optimal Growth Temperature (°C) | Average RGPD (mm/day) |
| --- | --- | --- | --- |
| Pc2113 | California | 22 | 7.34 ± 0.28 |
| 578 | California | 28 | 9.59 ± 1.29 |
| 647 | California | 28 | 6.35 ± 0.34 |
| FG 24-2 | California | 28 | 5.51 ± 0.17 |
| FG 29-1 | California | 28 | 5.97 ± 0.16 |
| FG 57-2 | California | 28 | 6.88 ± 0.28 |
| FG 90-2 | California | 28 | 8.25 ± 0.33 |
| Gary 1 | California | 28 | 9.85 ± 0.26 |
| Gary 2 | California | 28 | 10.23 ± 0.58 |
| AVR-001 | Florida | 28 | 5.14 ± 0.24 |
| AVR-008 | Florida | 30 | 4.95 ± 0.26 |
| F13-2 | Florida | 28 | 6.92 ± 0.14 |
| FL 2-1 | Florida | 28 | 9.59 ± 0.15 |
| FL 2-2 | Florida | 28 | 9.91 ± 0.12 |
| FL 4-1 | Florida | 30 | 8.30 ± 0.07 |
| R23 T3-4 | Florida | 30 | 8.16 ± 0.30 |
| R23 T3-5 | Florida | 30 | 7.85 ± 0.60 |
| R8 T1-1 | Florida | 28 | 8.28 ± 0.49 |
| HI-1-SZ1 | Hawaii | 28 | 8.52 ± 0.34 |
| HI-14-SZ1 | Hawaii | 28 | 6.73 ± 0.25 |
| HI-2-SZ1 | Hawaii | 28 | 8.42 ± 0.23 |

**Figure 1.**
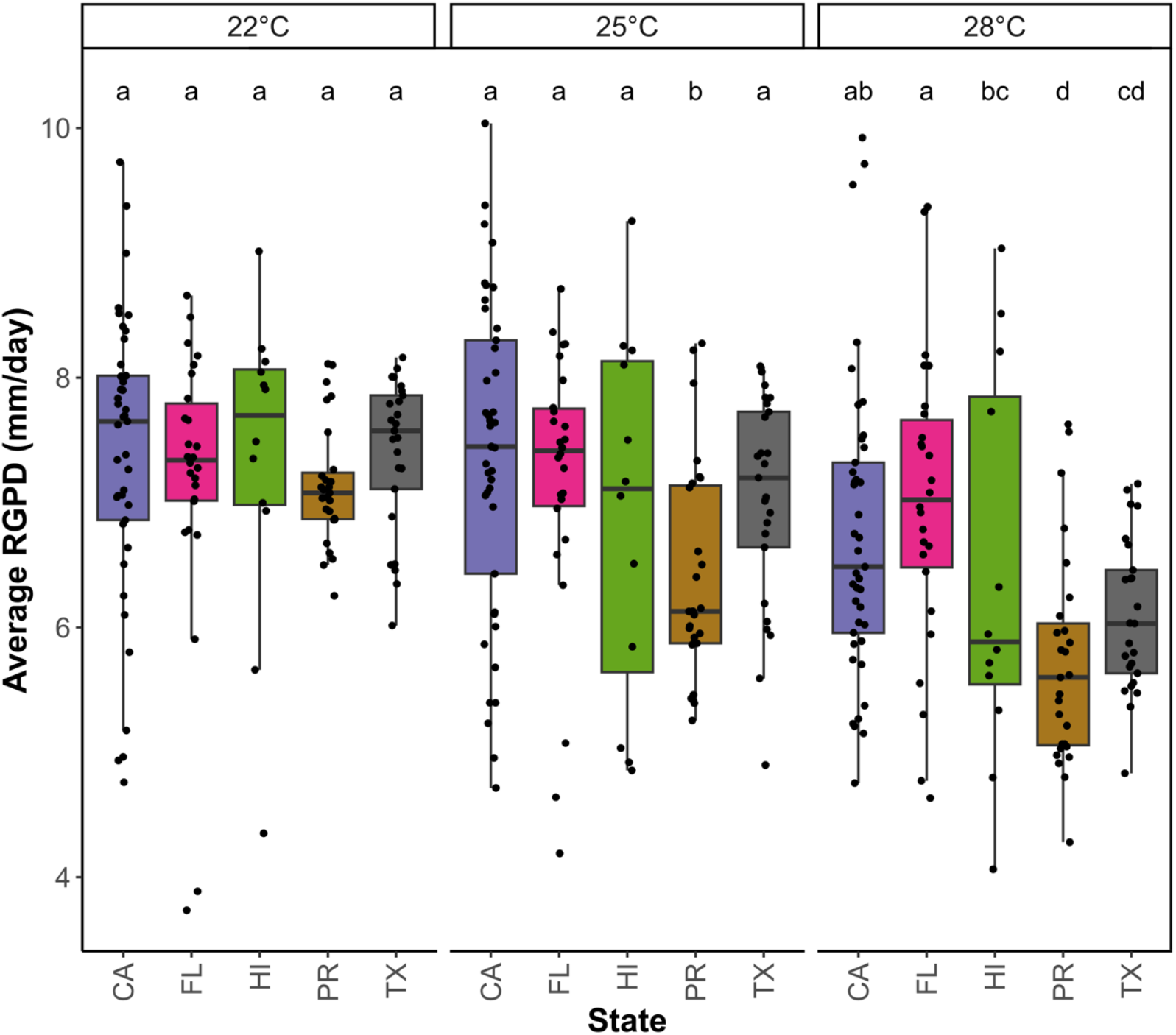
Phenotypic variability of *Phytophthora cinnamomi* isolates collected from California (CA), Florida (FL), Hawaii (HI), Puerto Rico (PR) and Texas (TX) regarding *in vitro* radial growth per day (RGPD) at 22°C, 25°C, and 28°C. Boxplots display the median RGPD and distribution of data of *P. cinnamomi* isolates at each temperature for each state. Bars followed by the same letter do not differ significantly in average RGPD between states within each temperature, according to the least squared means test at *P* < 0.05.

**Figure 2.**
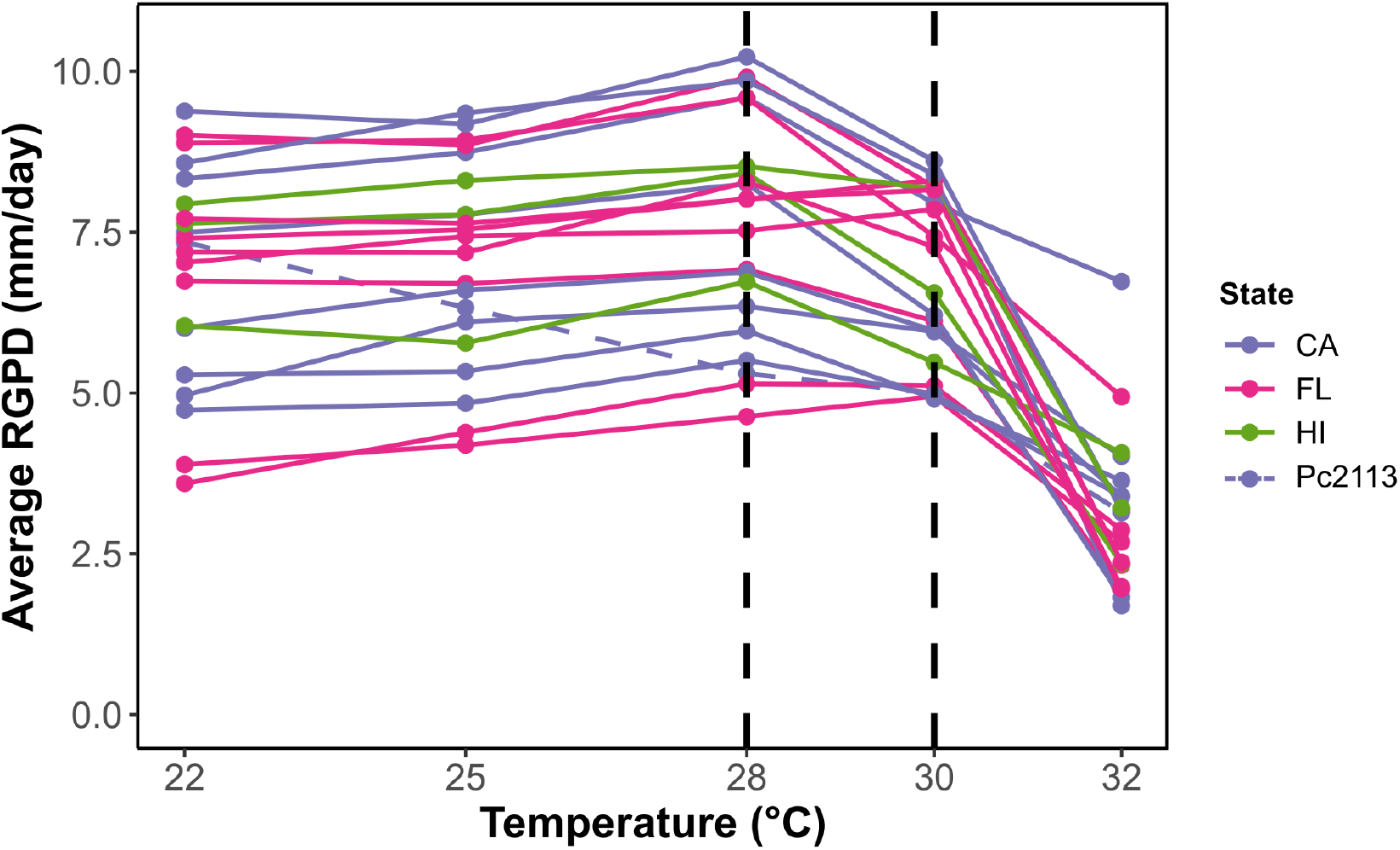
Assessment of *in vitro* radial growth per day (RGPD) and optimal growth temperature of the *Phytophthora cinnamomi* isolates exhibiting continued growth at 28°C. Isolates from California (CA), Florida (FL), and Hawaii (HI) were tested for their RGPD at 22°C, 25°C, 28°C, 30°C and 32°C. Optimal growth temperatures are indicated as dashed vertical black lines. Pc2113 isolate was used as internal control and is indicated as dashed purple line.

### 3.2 Isolates from California and Florida exhibit reduced sensitivity to potassium phosphite

All isolates were evaluated for their *in vitro* potassium phosphite sensitivity measured as EC_50_ (µg/ml) values which ranged from 4.61 (Gary 2 from CA) to 763.13 µg/ml (592 from CA). Significant differences were detected both within and among states (Figure 3; Supplementary Table S1). California isolates exhibited the greatest EC_50_ range, followed by isolates from Florida (10.55 to 238.51 µg/ml), Hawaii (5.93 to 76.04 µg/ml), Texas (10.17 to 33.67 µg/ml), and Puerto Rico (7.01 to 22.29 µg/ml) (Figure 3; Supplementary Table S1). These EC_50_ values were distributed into five bins with EC_50_ values <50 µg/ml (n = 100), 51-100 µg/ml (n = 11), 101-250 µg/ml (n = 5), 251-500 µg/ml (n = 5), and 501-800 µg/ml (n = 4). The frequency distribution of potassium phosphite EC_50_ values was unimodal and predominantly skewed toward high sensitivity (<50 µg/ml; n = 100) (Figure 3A). Average EC_50_ values for isolates from Puerto Rico, Texas and Hawaii (all <100 µg/ml) were significantly different from those of isolates from Florida and California (Figure 3B; Supplementary Table S3). Overall isolates from California and Florida showed reduced sensitivity (100 to 250 µg/ml) compared to other states, and nine California isolates exceeded 250 µg/ml, meeting the threshold for potassium phosphite resistance in this study (Figure 3; Supplementary Table S1). These patterns suggest a strong capacity for resistance development in regions where potassium phosphite is heavily used.

**Figure 3.**
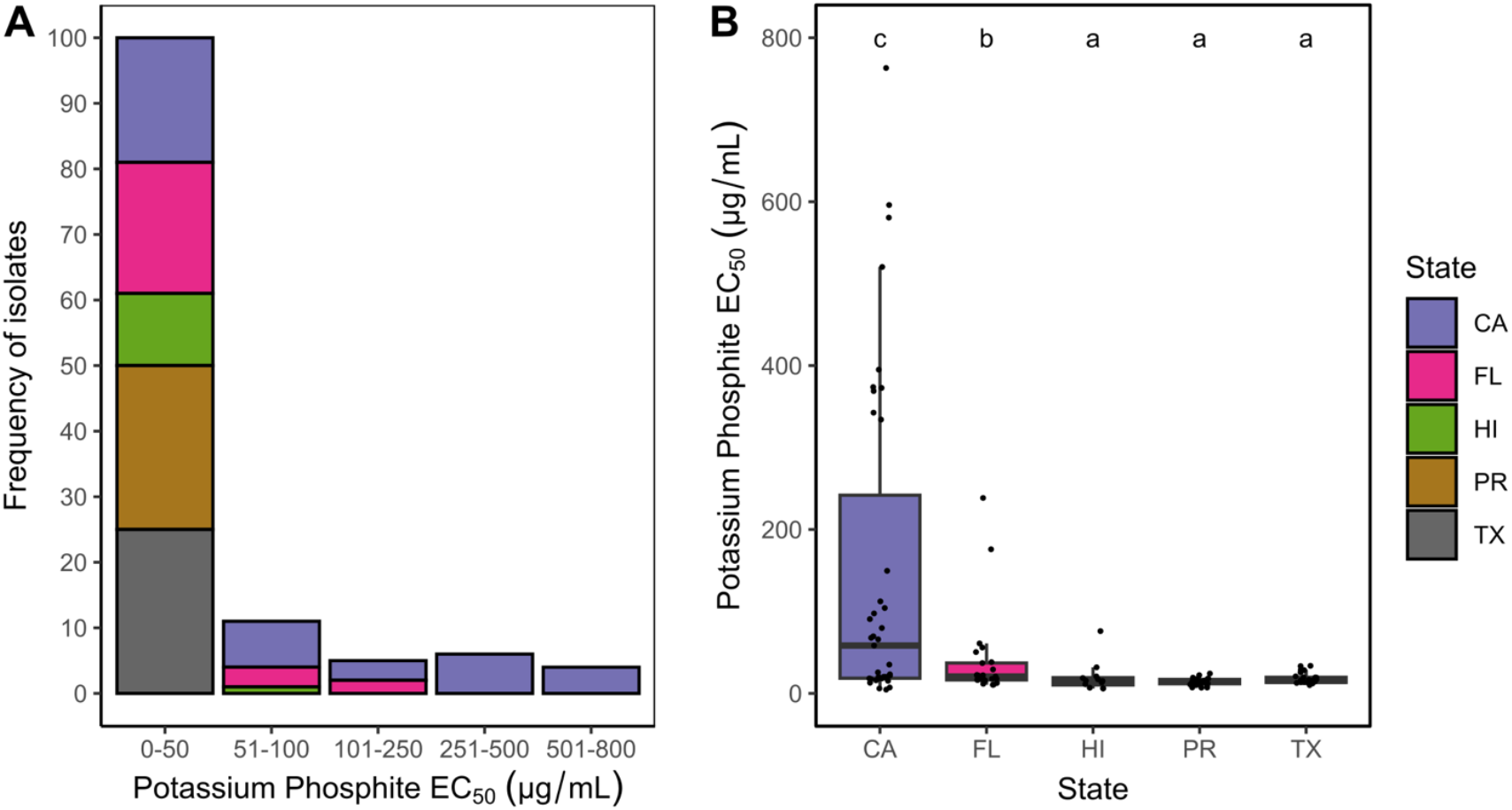
*In vitro* potassium phosphite sensitivity of *Phytophthora cinnamomi* isolates. **(A)** Frequency histograms of EC_50_ values (µg/ml) of 125 isolates. Bar heights indicate the number of isolates within each bin. Bin widths were assigned based on distributions of potassium phosphite sensitivities previously reported for *P. cinnamomi* isolates from avocado in California (Belisle et al., 2019b; Shands et al., 2025). **(B)** Boxplots showing the median and distribution of *in vitro P. cinnamomi* potassium phosphite EC_50_ values (µg/ml) for each state. Bars followed by the same letter did not significantly differ in the average EC_50_ value, according to the least squared means test at *P* < 0.05. Bars are color-coded based on the geographic origin of isolates (CA = California, FL = Florida, HI = Hawaii, PR = Puerto Rico, and TX = Texas).

### 3.3 Baseline sensitivities to Oomycota fungicides across U.S.A. avocado producing regions

To establish baseline sensitivity across states and enable comparison with established California benchmarks, all isolates were tested against five Oomycota fungicides: ethaboxam, mandipropamid, mefenoxam, fluopicolide, and oxathiapiprolin. All isolates were sensitive to the five Oomycota-targeting fungicides evaluated, although significant variation in EC_50_ values occurred within and among states. Across all isolates, EC_50_ values for ethaboxam ranged from 0.012 µg/ml (PR 5-1 from Puerto Rico) to 0.105 µg/ml (FG 57-2 from California), while mandipropamid EC_50_ values spanned from 0.0018 µg/ml (R8 T1-3 from Florida) to 0.0160 µg/ml (AVO 2-3 from Puerto Rico). For mefenoxam, EC_50_ values ranged between 0.011 µg/ml (LF 2-2 from Texas) and 0.174 µg/ml (AVO 2-3 from Puerto Rico). Fluopicolide displayed a broader range, with EC_50_ values from 0.08 µg/ml (R8 T1-3 from Florida) to 0.61 µg/ml (FG 31-2 from California), representing the least toxic of these five fungicides tested. Oxathiapiprolin exhibited the highest potency, with EC_50_ values from 0.00012 µg/ml (FG 62-1 from California) to 0.00064 µg/ml (AVO 2-3 from Puerto Rico) (Supplementary Table S1 and Supplementary Figure S2). EC_50_ values were grouped into Scott’s bins and displayed unimodal distributions with no evidence of reduced sensitivity or resistance among isolates from any of the avocado producing regions evaluated (Supplementary Figure S2).

Average EC_50_ values differed slightly among states, less than a 1-fold difference for each fungicide, though differences were statistically significant for all fungicides except mandipropamid (Figure 4; Supplementary Table S3). Across all states, average EC_50_ values remained below 0.036 ug/ml (ethaboxam), 0.007 µg/ml (mandipropamid), 0.055 µg/ml (mefenoxam), 0.37 µg/ml (fluopicolide), and 0.0004 µg/ml (oxathiapiprolin). Among the fungicides, fluopicolide was the least toxic, while oxathiapiprolin was the most potent (Figure 4; Supplementary Table S2).

**Figure 4.**
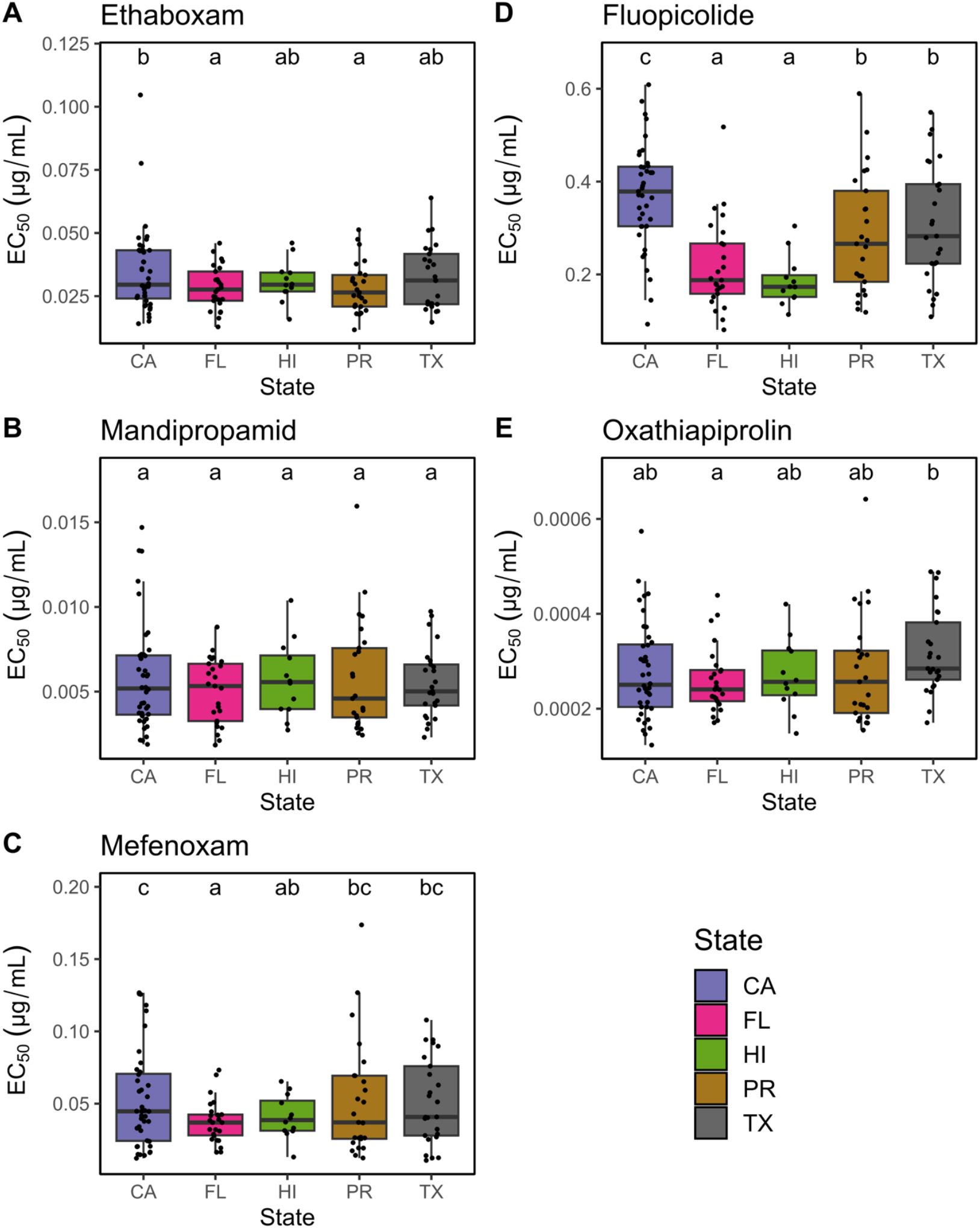
*In vitro* sensitivity of *Phytophthora cinnamomi* isolates to fungicides. Boxplots showing the median and distribution of the EC_50_ values (µg/ml) for ethaboxam **(A)**, mandipropamid **(B)**, mefenoxam **(C)**, fluopicolide **(D)**, and oxathiapiprolin **(E).**Bars followed by the same letter did not significantly differ in the average EC_50_ value, according to the least squared means test at *P* < 0.05. Bars are color-coded based on the geographic origin of isolates (CA = California, FL = Florida, HI = Hawaii, PR = Puerto Rico, and TX = Texas).

### 3.4 Extensive virulence variability observed among U.S. *P. cinnamomi* populations

Virulence was quantified as percent diseased roots in avocado seedlings and as AUDPC following inoculation of D’Anjou pear fruits. Uninoculated avocado controls remained mostly healthy (94.5%, *data not shown*). Inoculated seedlings exhibited significant differences in disease severity, ranging from 29% to 80% diseased roots, with California isolates spanning the widest range (30% to 80%). The most and least virulent isolates when infecting avocado were from California and Hawaii, respectively (Figure 5A; Supplementary Table S1). Hawaiian isolates exhibited significant lower virulence than isolates from California, Florida, and Puerto Rico. Consistent with previous findings (Shands et al., 2025), CA-South isolates remained significantly more virulent than CA-North isolates (55.6% vs. 45% diseased roots) on UC2001 seedlings (Supplementary Table S2).

**Figure 5.**
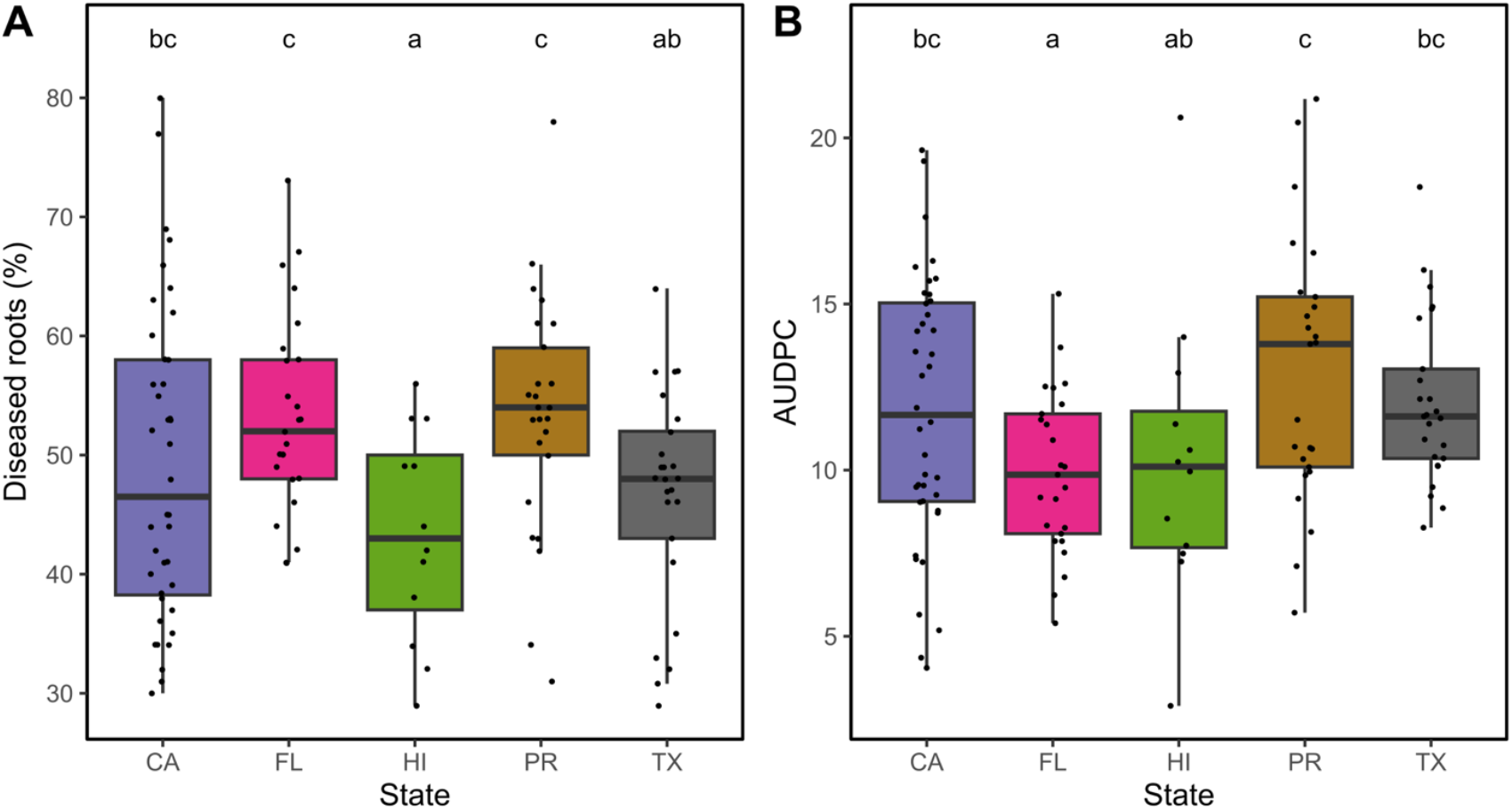
Virulence phenotype of *Phytophthora cinnamomi* isolates assessed in two different hosts. Boxplots showing the median and distribution of **(A)** percentage of diseased roots of UC2001 avocado seedling five-weeks post inoculation with *P. cinnamomi* isolates and **(B)** area under the disease progress curve (AUDPC) values of *P. cinnamomi* inoculated D’Anjou pears. Bars followed by the same letter did not show significant differences in virulence measurements according to the least squared means test at *P* < 0.05. Bars are color-coded based on the geographic origin of isolates (CA = California, FL = Florida, HI = Hawaii, PR = Puerto Rico, and TX = Texas).

Because *P. cinnamomi* can infect a wide range of plants, we also determined if the virulence of these isolates would translate across different hosts and tissue types. To address this, we inoculated D’Anjou fruits pears. We detected significant differences in the AUDPC values amongst all isolates, ranging from 2.9 (HI-5-SZ1) to 21.17 (AVO 1-1), with isolates from Hawaii exhibiting the largest variability (2.9-20.61) followed by those from Puerto Rico (5.71-21.17). On pear fruit, the most virulent isolates originated from Puerto Rico and the least from Hawaii (Figure 5B; Supplementary Table S1). Isolates from Hawaii differed significantly in AUDPC values when compared to those collected from California, Puerto Rico, and Texas, and unlike the avocado assay, California and Florida isolates differed significantly in their virulence when using pear fruits (Figure 5B; Supplementary Table S3). In contrast to the avocado assay, no significant differences were detected between CA-South and CA-North isolates on pear fruits (Supplementary Figure S2).

## 4 Discussion

We previously documented substantial phenotypic variability in *P. cinnamomi* populations associated with avocado PRR in California (Belisle et al., 2019a; Belisle et al., 2019b; Shands et al., 2025). In this study, we expanded that work by characterizing pathogen populations from Florida, Puerto Rico, Hawaii, Texas, and the most current California population for optimal growth temperature, radial growth rate, *in vitro* fungicide sensitivity, and virulence on both D’Anjou pear fruit and moderately resistant UC2001 avocado seedlings. Phenotypic studies of *P. cinnamomi* affecting avocado in the U.S.A. have been largely restricted to California (Belisle et al., 2019a; Belisle et al., 2019b; Pagliaccia et al., 2013; Shands et al. 2025), while research in other states has focused on pathogen detection and screening for rootstock resistance (Abdulridha et al., 2019; Crandall, 1948; Hwang, 1978; Ploetz et al., 2001; Zentmyer, 1976). Thus, this work represents the first comparative phenotypic characterization of *P. cinnamomi* populations across multiple avocado producing states in the U.S.A. and established baseline *in vitro* sensitivities for three registered and three unregistered Oomycota fungicides (Hoyt et al., 2026).

Although *Phytophthora cinnamomi* is heterothallic (A1 and A2 mating types), only clonal A2 populations have been reported in avocado orchards around the world (Dobrowolski et al., 2003; Engelbrecht et al., 2022; Ochoa-Fuentes et al., 2007; Pagliaccia et al., 2013; Shakya et al., 2021; Shands et al., 2025). Shakya et al. (2021) identified two global A2 panglobal clonal lineages (PcG1-A2 and PcG2-A2) and Pagliaccia et al. (2013) described two clonal A2 clades in California (A2 clade I and A2 clade II). Despite the absence of sexual recombination, extensive phenotypic differentiation among clonal populations has been repeatedly documented across hosts and regions (Belisle et al., 2019a; Belisle et al., 2019b; Calle-Henao et al., 2020; Hüberli et al., 2001; Ochoa-Fuentes et al., 2015; Shands et al., 2025). The ability to generate large phenotypic variability without sexual reproduction and successfully adapt to climate and control methods highlights the plasticity of this pathogen and demonstrates the challenges for developing effective and durable avocado PRR control methods (Belisle et al., 2019a; Belisle et al., 2019b; Calle-Henao et al., 2020; Hüberli et al., 2001; Shands et al., 2025). Consistent with previous observations, in this study, we also detected broad phenotypic diversity for key pathogen traits among isolates from all states evaluated.

Optimal temperature studies from avocado and other hosts have typically reported *P. cinnamomi* optimal growth between 24-28°C in California (Belisle et al., 2019a; Shands et al., 2025), Mexico (Mondragón-Flores et al., 2022), Australia (Hüberli et al., 2001), and the Pacific Northwest region of the United States (Scagel et al., 2023). In this study, most isolates grew best at 22°C or 25°C, with reduced growth at 28°C. Isolates from Puerto Rico and Texas were particularly adapted to 22°C, consistent with cooler local climates in collection sites. Similar adaptation to cooler environments has been reported in sub-alpine *P. cinnamomi* populations infecting eucalyptus and avocado (Khaliq et al., 2020). Conversely, we identified isolates from California, Florida, and Hawaii with optimal growth temperatures of 28 or 30°C (Table 2), supporting adaptation to warmer climates. As reported previously (Shands et al., 2025), isolates from CA-South were better adapted to higher temperatures than CA-North isolates, supporting persistent local thermal selection in this state. Higher optimal growth temperatures (>30°C) have also been reported in other regions (Khaliq et al., 2020). These findings align with the hypothesis that fungal plant pathogens can evolve to acquire thermal stability in response to regional climates (Miñana-Posada et al., 2025), suggesting similar local adaptation within clonal *P. cinnamomi* lineages associated with avocado PRR in U.S.A.

Excluding potassium phosphite, all Oomycota fungicides exhibited unimodal EC_50_ distributions with narrow ranges across isolates, indicating high baseline sensitivity and low current risk of resistance selection when used appropriately and in rotation programs. Avocado growers rely upon several registered chemicals to control PRR, including phosphonate-based products (e.g., potassium phosphite), phenylamides (e.g., mefenoxam), and only in California, the recently registered piperidinyl thiazole isoxazoline fungicide, oxathiapiprolin (Adaskaveg et al., 2024; Dann and Gazis, 2024). Previous California studies established baseline sensitivities for these compounds and several additional Oomycota fungicides (Belisle et al., 2019b). Because comparable data were lacking for other U.S.A. states, our results provide the first baseline sensitivities for isolates from Florida, Puerto Rico, Texas, and Hawaii, creating a benchmark for monitoring the emergence or introductions of fungicide-resistant populations.

With the exception of potassium phosphite, the baseline sensitivities of *P. cinnamomi* populations to these fungicides in other states were similar to those previously reported in California, reflecting natural sensitivity levels of this pathogen. As in earlier studies (Belisle et al., 2019a; Belisle et al., 2019b), oxathiapiprolin was the most effective fungicide, exhibiting the lowest EC_50_ values. This fungicide also has been shown to be highly toxic to other *Phytophthora* spp. with inhibitory values 10-to 1000-fold lower than those reported for ethaboxam, fluopicolide, mandipropamid, and mefenoxam (Ji and Csinos, 2015; Miao et al., 2016; Qu et al., 2016; Riley et al., 2024). Mefenoxam and metalaxyl have been commonly used to control *P. cinnamomi*, especially in nurseries and natural ecosystems (Dunstan et al., 2008; Hu et al., 2010). Mefenoxam EC_50_ values were consistent with studies in avocado and ornamentals (Belisle et al., 2019a; Belisle et al., 2019b; Benson and Grand, 2000; Duan et al., 2008; Hu et al., 2010), reflecting continued effectiveness when used properly.

In contrast, elevated potassium phosphite EC_50_ values, particularly among California and Florida isolates, mirror increasing reports of potassium phosphite-resistant *P. cinnamomi* isolates in California (Adaskaveg et al., 2024; Shands et al., 2025), Australia (Andronis et al., 2024), South Africa (Ma and McLeod, 2014), and New Zealand (Hunter et al., 2023). Similar resistance to this chemical has been documented in *P. cinnamomi* isolates infecting other hosts (Vegh et al., 1985) and in other oomycete pathogens (Brown et al., 2004; Hao et al., 2021). Although most isolates remained sensitive (<50 µg/ml), significantly higher EC_50_ values were concentrated in regions with long histories of potassium phosphite applications such as California and Florida (Belisle et al., 2019b; Cohen and Coffey, 1986; Mossler and Crane, 2008; Shands et al., 2025). Notably, we report for the first time isolates from Santa Barbara with EC_50_ values as high as 763.13 µg/ml. This result is unexpected given previous findings that CA-North isolates (Santa Barbara and Ventura) were more sensitive than CA-South populations (Belisle et al., 2019a; Belisle et al., 2019b; Shands et al., 2025). This shift may reflect movement of less-sensitive isolates from CA-South to CA-North which is consistent with the recent detection of clonal A2 clade II isolates in CA-North, formerly restricted to just CA-South (Santa Barbara) (Pagliaccia et al., 2013; Shands et al., 2025).

Significant variability in virulence was also detected in this study in avocado and pear fruits. Virulence variability has been reported in *P. cinnamomi* across hosts and regions (Ochoa-Fuentes et al., 2015; Vitale et al., 2018; Robin & Desprez-Loustau, 1998; Serrano & Garbelotto, 2020). Moreover, clonal populations of *P. cinnamomi* associated with avocado PRR in Southern California growing regions have been reported to be more virulent in moderately resistant PRR rootstocks including Dusa^®^ and UC2001 (Belisle et al., 2019a; Belisle et al., 2019b; Hoyt et al., 2026; Shands et al., 2025). Studies assessing the reduction in resistance of other avocado cultivars towards *P. cinnamomi* isolates have also been reported in Mexico (Sánchez-González et al., 2019). As observed previously, virulence patterns often differed between hosts with several isolates exhibiting opposite virulence rankings in pear and avocado. Such host-specific differences are consistent with reports that infection biology and transcriptomic responses of *P. cinnamomi* vary depending on the host tissue and species (Shands et al., 2024; Shands et al., 2025). Thus, in our study, the virulence of the pathogen populations from avocado orchards around the U.S.A. were characterized using different infection assays in D’Anjou pear fruits and avocado UC2001 seedlings. Pear fruits have been used to assess pathogenicity and as bait for different *Phytophthora* spp., including *P. cinnamomi* (Sanchez et al., 2019; Shands et al., 2025; Swiecki et al., 2024). The D’Anjou pear fruit virulence assay revealed wide variability across isolates, with Texas, Puerto Rico, and California isolates exhibiting the highest severity. Shands et al. (2025) reported that CA-North isolates were more virulent than CA-South isolates when pear fruits were used to evaluate their virulence and these results were completely opposite when virulence was tested in avocado seedlings. Here, we did not detect significant differences between isolates collected from these two growing regions (Supplementary Table S2), however, we found several isolates including Pc2113, Pc2109, and FG 90-2 from California, R23 T3-4 from Florida, and TL 5-2 from Texas whose virulence phenotypes were opposite when tested in these two hosts (Supplementary Table S1).

Virulence variability among *P. cinnamomi* isolates associated with avocado PRR have been reported in Mexico and California. Ochoa-Fuentes et al. (2015) found that isolates from Mexico were more virulent in Peribán and Uruapan compared to those from Salvador Escalante. Consistent with our prior work (Belisle et al., 2019a; Shands et al., 2025), the CA-South isolates in this study remained more virulent on UC2001 seedlings than CA-North isolates. In addition, isolates from California, Florida, and Puerto Rico were generally more virulent on avocado than those from Hawaii. The most virulent isolate was again collected from San Diego County, aligning with previous reports of heightened virulence in Southern California populations (Belisle et al., 2019a; Shands et al., 2025). Identifying the most virulent *P. cinnamomi* isolates among domestic avocado orchards will aid in creating effective PRR management strategies, including the screening of avocado germplasm to develop rootstocks harboring resistance to these virulent populations and the development of phenotype-specific diagnostic tools that could monitor the introduction and spread of more virulent isolates among these avocado producing regions.

In summary, we detected extensive phenotypic variability within clonal *P. cinnamomi* populations across major U.S.A. avocado producing regions and documented the presence of highly virulent, potassium phosphite-resistant, and thermally adapted isolates. Given the pathogen’s wide host range (>5000 species), virulence assessments across multiple hosts are valuable for both agricultural and natural diversity. These findings have important implications for PRR management and establish a critical foundation for future work linking phenotypic patterns to the genetic diversity of U.S.A. *P. cinnamomi* populations. Understanding the molecular basis of this pathogen’s phenotypic plasticity and adaptability will be essential for developing sustainable, effective, and durable PRR control strategies to support the competitiveness and long-term viability of the U.S.A. avocado industry.

## 6 Author Contributions

Conceptualization and funding acquisition: P.M, J.C., R.G., L.C., M.T., J.J., R.G., and J.A. Experimental design: P.M. and B.H.

Research Investigation: B.H. and S.S.

Resources: B.H., J.C., M.N., R.G., L.C., A.A., M.T., R.G., L.S., and J.A.

Data analyses: B.H.

Writing, Review & Editing: P.M. and B.H. All authors reviewed and approved the final manuscript.

## 8 Acknowledgments

We thank Matthew Elvena and Amber Newsome for collecting root and soil samples throughout California avocado orchards; Jocelyn Leos and Vanessa Hua for their assistance in phenotypic assays; Dr. Nathan Riley, Dr. Chaydon O’Fallon, and Dr. Albert Nguyen for their technical guidance regarding SGP assays; Dr. Aidan Shands for his technical guidance regarding phenotypic assessment assays; and Dr. Greg Browne for providing us with *P. cinnamomi* isolates from California walnut trees.

## 12 Supplementary Material

**Supplementary Figure S1.**
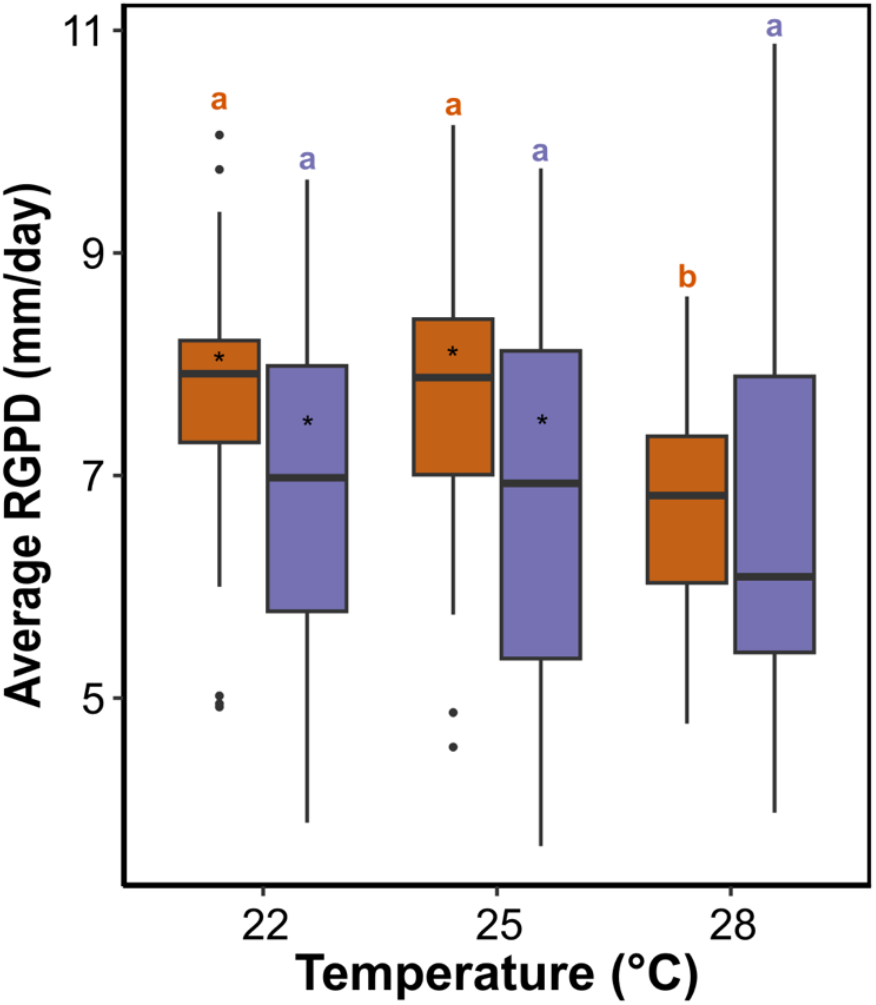
Boxplots showing the average *in vitro* radial growth per day (RGPD) of *P. cinnamomi* isolates collected from CA–North (orange) and CA–South (purple) at 22°C, 25°C, and 28°C. Letters indicate significant differences across growth temperatures within each avocado growing regions and asterisks (*) indicate significant differences detected for isolates between growing regions at each temperature assayed. Significant differences were calculated using GLMM analyses followed by least squared means test at *P* < 0.05.

**Supplementary Figure S2.**
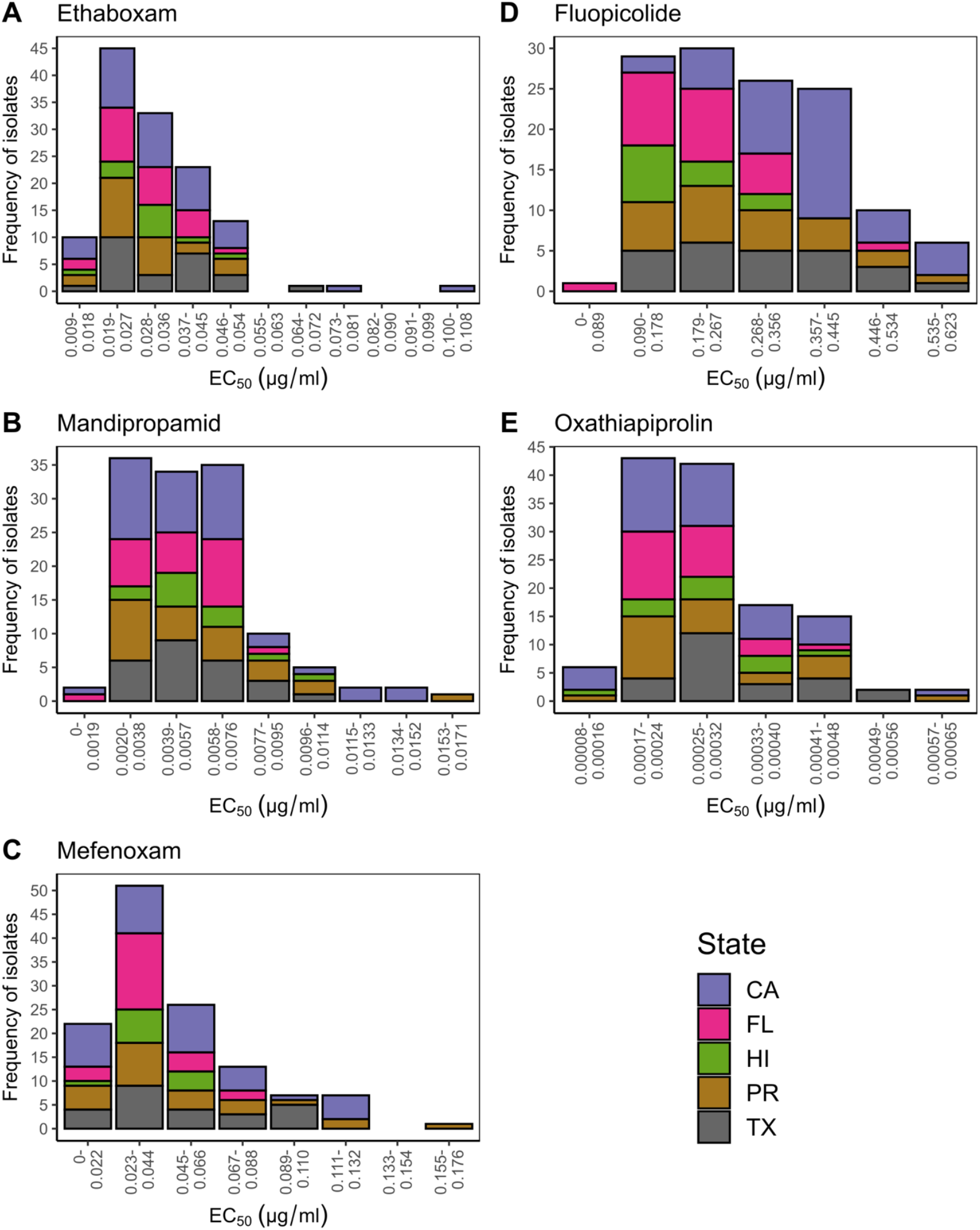
Frequency histograms of *in vitro* fungicide sensitivities (µg/ml) of *P. cinnamomi* isolates to ethaboxam **(A)**, mandipropamid **(B)**, mefenoxam **(C)**, fluopicolide **(D)**, and oxathiapiprolin **(E)** collected from different avocado producing states in the U.S.A. Bar heights indicate the number of isolates within each bin. Bin widths were calculated based on Scott’s rule (1979). Bars are color-coded based on the geographic origin of isolates (CA = California, FL = Florida, HI = Hawaii, PR = Puerto Rico, and TX = Texas).

**Supplementary Table S1**. Phenotypic variation of *in vitro* radial growth rate per day (RGPD), *in vitro* fungicide sensitivity, and virulence of all *Phytophthora cinnamomi* isolates. The average *in vitro* radial growth rate per day (RGPD) at 22°C, 25°C, 28°C, 30°C and 32°C, the average *in vitro* EC_50_ values (µg/ml) of potassium phosphite, mefenoxam, oxathiapiprolin, ethaboxam, fluopicolide, and mandipropamid and the average virulence in D’Anjou pears (AUDPC) and UC2001 avocado seedlings (% diseased roots) are displayed for each isolate. Values followed by the same letter are not significantly different according to the least squared means test at *P* < 0.05.

**Supplementary Table S2**. *In vitro* radial growth rate per day (RGPD) and virulence of *Phytophthora cinnamomi* isolates collected from CA-North and CA-South avocado growing regions. Average *in vitro* RGPD at 22°C, 25°C, and 28°C. Average virulence in D’Anjou pears (AUDPC) and UC2001 avocado seedlings (% diseased roots) are also shown. Values followed by the same letter are not significantly different according to the least squared means test at *P* < 0.05.

**Supplementary Table S3**. Phenotypic trait variability of *Phytophthora cinnamomi* isolates by state (California, Florida, Hawaii, Texas, and Puerto Rico). *In vitro* radial growth rate per day (RGPD) at 22°C, 25°C, 28°C, 30°C and 32°C, EC_50_ values (µg/ml) of each fungicide, and virulence assessed in D’Anjou pears as AUDPC) and UC2001 avocado seedlings as % diseased roots are indicated. Values followed by the same letter are not significantly different according to the least squared means test at *P* < 0.05.

## References

Abdulridha, J., Ehsani, R., Abd-Elrahman, A., and Ampatzidis, Y. (2019). A remote sensing technique for detecting laurel wilt disease in avocado in presence of other biotic and abiotic stresses. Comput. Electron. Agric. 156, 549–557. doi: 10.1016/j.compag.2018.12.018

Adaskaveg, J. E., Hao, W., and Förster, H. (2015). Postharvest strategies for managing Phytophthora brown rot of citrus using potassium phosphite in combination with heat treatments. Plant Dis. 99, 1477–1482. doi: 10.1094/PDIS-01-15-0040-RE

Adaskaveg, J. E., Förster, H., and O’Fallon, C. (2024). New fungicides for managing Phytophthora diseases of tree crops with foliar and soil applications. J. Plant Dis. Prot. 131, 1203–1209. doi: 10.1007/s41348-024-00873-6

Andronis, C. E., Jacques, S., Lopez-Ruiz, F. J., Lipscombe, R., and Tan, K.-C. (2024). Proteomic analysis revealed that the oomyceticide phosphite exhibits multi-modal action in an oomycete pathosystem. J. Proteomics 301, 105181. doi: 10.1016/j.jprot.2024.105181

Bates, D., Mächler, M., Bolker, B., and Walker, S. (2015). Fitting linear mixed-effects models using lme4. J. Stat. Softw. 67, 1–48. doi: 10.18637/jss.v067.i01

Belisle, R. J., McKee, B., Hao, W., Crowley, M., Arpaia, M. L., Miles, T. D., et al. (2019a). Phenotypic characterization of genetically distinct Phytophthora cinnamomi isolates from avocado. Phytopathology 109, 384–394. doi: 10.1094/PHYTO-09-17-0326-R

Belisle, R. J., Hao, W., McKee, B., Arpaia, M. L., Manosalva, P., and Adaskaveg, J. E. (2019b). New Oomycota fungicides with activity against Phytophthora cinnamomi and their potential use for managing avocado root rot in California. Plant Dis. 103, 2024–2032. doi: 10.1094/PDIS-09-18-1698-RE

Bender, G. S., Menge, J. A., and Arpaia, M. L. (2012). “Avocado rootstocks” in Avocado Production in California: A Cultural Handbook for Growers, Book 1: Background Information, ed. G. S. Bender (Oakland, CA: University of California Agriculture and Natural Resources), 20–30.

Benson, D. M., and Grand, L. F. (2000). Incidence of Phytophthora root rot of Fraser fir in North Carolina and sensitivity of isolates of Phytophthora cinnamomi to metalaxyl. Plant Dis. 84, 661–664. doi: 10.1094/PDIS.2000.84.6.661

Bilodeau, G. J., Martin, F. N., Coffey, M. D., and Blomquist, C. L. (2014). Development of a multiplex assay for genus- and species-specific detection of Phytophthora based on differences in mitochondrial gene order. Phytopathology 104, 733–748. doi: 10.1094/PHYTO-09-13-0263-R

Brown, S., Koike, S. T., Ochoa, O. E., Laemmlen, F., and Michelmore, R. W. (2004). Insensitivity to the fungicide fosetyl-aluminum in California isolates of the lettuce downy mildew pathogen, Bremia lactucae. Plant Dis. 88, 502–508. doi: 10.1094/PDIS.2004.88.5.502

Browne, G. T., Leslie, C. A., Grant, J. A., Bhat, R. G., Schmidt, L. S., Hackett, W. P., et al. (2015). Resistance to species of Phytophthora identified among clones of Juglans microcarpa × J. regia. HortScience 50(8), 1136–1142. doi: 10.21273/HORTSCI.50.8.1136

Burgess, T. I., Scott, J. K., Mcdougall, K. L., Stukely, M. J. C., Crane, C., Dunstan, W. A., et al. (2017). Current and projected global distribution of Phytophthora cinnamomi, one of the world’s worst plant pathogens. Global Change Biol. 23, 1661–1674. doi: 10.1111/gcb.13492

Calle-Henao, C., Gonzales-Jaimes, E. P., Arango-Isaza, R. E., and Saldamando-Benjumea, C. I. (2020). Isolation and identification of Phytophthora cinnamomi collected in avocado (Persea americana) from Northeast Colombia. Trop. plant pathol. 45, 402–414. doi: 10.1007/s40858-020-00337-w

Coffey, M. D. (1987). Phytophthora root rot of avocado: an integrated approach to control in California. Plant Dis. 71, 1046–1053.

Cohen, Y., and Coffey, M. D. (1986). Systemic Fungicides and the Control of Oomycetes. Annu. Rev. Phytopathol. 24, 311–338. doi: 10.1146/annurev.py.24.090186.001523

Crandall, B. S. (1948). “Phytophthora cinnamomi root rot of avocados under tropical conditions,” in California Avocado Soc. 1948 Yearb. 33, 76–81.

Crane, J. H., Manosalva, P., Faber, B. A., Barrientos-Priego, A. F., Focht, E. D., Arpaia, M. L., Bender, G. S., and Balerdi, C. F. (2024). “Cultivars and rootstocks” in The Avocado: Botany, Production and Uses, 3rd ed., eds. B. Schaffer, B. N. Wolstenholme, and A. W. Whiley (Wallingford, United Kingdom: CABI), 212–263.

Dann, E.K., Gazis, R. (2024). “Foliar, Fruit and Soilborne Diseases” in The Avocado: Botany, Production and Uses, 3rd ed., eds. B. Schaffer, B. N. Wolstenholme, and A. W. Whiley (Wallingford, United Kingdom: CABI), 212–263.

Dobrowolski, M. P., Tommerup, I. C., Shearer, B. L., and O’Brien, P. A. (2003). Three clonal lineages of Phytophthora cinnamomi in Australia revealed by microsatellites. Phytopathology 93, 695–704. doi: 10.1094/PHYTO.2003.93.6.695

Dobrowolski, M. P., Shearer, B. L., Colquhoun, I. J., O’Brien, P. A., and Hardy, G. E. StJ. (2008). Selection for decreased sensitivity to phosphite in Phytophthora cinnamomi with prolonged use of fungicide. Plant Pathol. 57, 928–936. doi: 10.1111/j.1365-3059.2008.01883.x

Dreistadt, S. H. (2007). “Avocado root rot or Phytophthora root rot” in Integrated Pest Management for Avocados, UC ANR Publication 3503 (Oakland, CA: University of California Agriculture and Natural Resources), 60–65.

Duan, C.-H., Riley, M. B., and Jeffers, S. N. (2008). Characterization of Phytophthora cinnamomi populations from ornamental plants in South Carolina, USA. Arch. Phytopathol. Plant Prot. 41, 14– 30. doi: 10.1080/03235400600628054

Dreher, M. L., and Davenport, A. J. (2013). Hass avocado composition and potential health effects. Crit. Rev. Food Sci. Nutr. 53, 738–750. doi: 10.1080/10408398.2011.556759

Dunstan, W. A., Rudman, T., Shearer, B. L., Moore, N. A., Dell, B., Crane, C., Barrett, S. and Hardy, G. E. St. J. (2008). “Containment and eradication of Phytophthora cinnamomi in native vegetation in South-Western Australia and Tasmania” in Proceedings of the Fourth Meeting of the IUFRO Working Party S07. 02. 09, Phytophthoras in Forests and Natural Ecosystems, eds. M. Garbelotto and S. J. Frankel (Albany, CA: U.S. Department of Agriculture, Forest Service, Pacific Southwest Research Station General Technical Report PSW-GTR-211), pp. 218–226.

Engelbrecht, J., Duong, T. A., Paap, T., Hubert, J. M., Hanneman, J. J., and van den Berg, N. (2022). Population genetic analyses of Phytophthora cinnamomi reveals three lineages and movement between natural vegetation and avocado orchards in South Africa. Phytopathology 112, 1568–1574. doi: 10.1094/PHYTO-10-21-0414-R

FAO. (2025). FAOSTAT. Avocado production statistics. Availaible online at: https://www.fao.org/faostat/en/#data/QCL/visualize (Accessed December 17, 2025).

Faber, B.A., Bender, G., and Eskalen, A. (2012) “Phytophthora root rot” in Avocado Production in California: A Cultural Handbook for Growers, Book 2: Integrated Pest Management for Avocado, ed. G. S. Bender (Oakland, CA: University of California Agriculture and Natural Resources), 61–65.

Hammami, A. M., Huang, K.-M., and Guan, Z. (2024). An overview of the avocado market in the United States. EDIS 2024. doi: 10.32473/edis-fe1150-2024

Hardham, A. R., and Blackman, L. M. (2018). Phytophthora cinnamomi. Mol. Plant Pathol. 19, 260– 285. doi: 10.1111/mpp.12568

Hao, W., Miles, T. D., Martin, F. N., Browne, G. T., Förster, H., and Adaskaveg, J. E. (2018). Temporal occurrence and niche preferences of Phytophthora spp. causing brown rot of citrus in the Central Valley of California. Phytopathology 108, 384–391. doi: 10.1094/PHYTO-09-17-0315-R

Hao, W., Förster, H., and Adaskaveg, J. E. (2021). Resistance to potassium phosphite in Phytophthora species causing citrus brown rot and integrated practices for management of resistant isolates. Plant Dis. 105, 972–977. doi: 10.1094/PDIS-06-20-1414-RE

Hawaii Department of Agriculture. (2024). Avocado statistics: State of Hawaii, 2017–2022. https://dab.hawaii.gov/add/files/2024/01/Avocado-Stats-2022_SOH_01.31.2024.pdf [Accessed December 17, 2025].

Hoyt, B. K., Mamo, B. E., Elvena, M., Newsome, A., Salas, S., Leos, J., Hua, V., Adaskaveg, J. E., and Manosalva, P. (2026). Reducing losses to Phytophthora cinnamomi by combining Oomycota fungicides and avocado rootstocks with different levels of resistance. Plant Dis. doi: 10.1094/PDIS-12-25-2484-RE

Hu, J., Hong, C., Stromberg, E. L., and Moorman, G. W. (2010). Mefenoxam sensitivity in Phytophthora cinnamomi Isolates. Plant Dis. 94, 39–44. doi: 10.1094/PDIS-94-1-0039

Hüberli, D., Tommerup, I. C., Dobrowolski, M. P., Calver, M. C., and Hardy, G. E. (2001). Phenotypic variation in a clonal lineage of two Phytophthora cinnamomi populations from Western Australia. Mycol. Res. 105, 1053–1064. doi: 10.1016/S0953-7562(08)61967-X

Hunter, S., McDougal, R., Williams, N., and Scott, P. (2023). Evidence of phosphite tolerance in Phytophthora cinnamomi from New Zealand avocado orchards. Plant Dis. 107, 393–400. doi: 10.1094/PDIS-05-22-1269-RE

Hwang, S. C. (1978). Biology of chlamydospores, sporangia, and zoospores of Phytophthora cinnamomi in soil. Phytopathology 68, 726. doi: 10.1094/Phyto-68-726

Ji, P., and Csinos, A. S. (2015). Effect of oxathiapiprolin on asexual life stages of Phytophthora capsici and disease development on vegetables. Ann. Appl. Biol. 166, 229–235. doi: 10.1111/aab.12176

Jung, T., Colquhoun, I. J., and Hardy, G. E. St. J. (2013). New insights into the survival strategy of the invasive soilborne pathogen Phytophthora cinnamomi in different natural ecosystems in Western Australia. For. Path. 43, 266–288. doi: 10.1111/efp.12025

Kavroulakis, N., Tziros, G. T., Mikalef, L., and Malandrakis, A. A. (2024). First report of Phytophthora cinnamomi causing root rot of avocado trees in Greece. Plant Dis. 108, 3185. doi: 10.1094/PDIS-05-24-0939-PDN

Khaliq, I., Hardy, G. E. St. J., and Burgess, T. I. (2020). Phytophthora cinnamomi exhibits phenotypic plasticity in response to cold temperatures. Mycol. Prog. 19, 405–415. doi: 10.1007/s11557-020-01578-4

Ko, W.-H. (2003). Long-term storage and survival structure of three species of Phytophthora in water. J. Gen. Plant. Pathol. 69, 186–188. doi: 10.1007/s10327-003-0033-3

Köhne, J. S. (1998). “Avocado tree spacing trends and size control in South Africa,” in Seminario Internacional de Paltos, Viña del Mar, Chile, 1998.

Kunta, M., da Graça, J., & Skaria, M. (2007). Molecular detection and prevalence of citrus viroids in Texas. HortScience, 42(3), 600–604. doi:10.21273/HORTSCI.42.3.600

Kurbetli, İ., Sülü, G., Aydoğdu, M., Woodward, S., and Bayram, S. (2020). Outbreak of Phytophthora cinnamomi causing severe decline of avocado trees in Southern Turkey. J. Phytopathol. 168, 533–541. doi: 10.1111/jph.12931

Lenth, R. V. (2016). Least-squares means: the R package lsmeans. J. Stat. Soft. 69(1), 1–33. 10.18637/jss.v069.i01

Lonsdale, J. H., Botha, T., and Kotzé, J. M. (1988). Preliminary trials to assess the resistance of three clonal avocado rootstocks to crown canker caused by Phytophthora cinnamomi. South African Avocado Growers’ Association Yearbook 11, 35–37.

López-Herrera, C. J., and Pérez-Jiménez, R. M. (1995). Morphology of Phytophthora cinnamomi isolates from avocado orchards in the coastal area of Southern Spain. J. Phytopathol. 143, 735–737. doi: 10.1111/j.1439-0434.1995.tb00232.x

Lowe, S., Browne, M., Boudjelas, S., and De Poorter, M. (2000). 100 of the World’s Worst Invasive Alien Species: A selection from the Global Invasive Species Database. Invasive Species Specialist Group.

Ma, J., and McLeod, A. (2014). In vitro sensitivity of South African Phytophthora cinnamomi isolates to phosphite at different phosphate concentrations. South African Avocado Growers’ Association Yearbook 37, 79–83.

Madhu, G. S., Rani, A. T., Muralidhara, B. M., Deepak, G. N., Rajendiran, S., Ayyandurai, M., et al. (2025). Multigene phylogeny and diversity of Phytophthora and Phytopythium species associated with avocado root rot in India and development of a point-of-care LAMP assay for Phytophthora cinnamomi and Phytopythium vexans. Physiol. Mol. Plant Pathol. 138, 102735. doi: 10.1016/j.pmpp.2025.102735

Menge, J. A., McKee, B. S., Guillmet, F. B., and Pond, E. C. (1999). Results of recent tests for root rot tolerance in avocado rootstocks in California. California Avocado Society Yearbook 83, 75–86.

Miao, J., Dong, X., Lin, D., Wang, Q., Liu, P., Chen, F., et al. (2016). Activity of the novel fungicide oxathiapiprolin against plant-pathogenic oomycetes. Pest Manag. Sci. 72, 1572–1577. doi: 10.1002/ps.4189

Miles, T. D., Martin, F. N., Robideau, G. P., Bilodeau, G. J., and Coffey, M. D. (2017). Systematic development of Phytophthora species-specific mitochondrial diagnostic markers for economically important members of the genus. Plant Dis. 101, 1162–1170. doi: 10.1094/PDIS-09-16-1224-RE

Miñana-Posada, S., Lorrain, C., McDonald, B. A., and Feurtey, A. (2025). Thermal adaptation in worldwide collections of a major fungal pathogen. MPMI 38, 252–264. doi: 10.1094/MPMI-09-24-0112-FI

Mondragón-Flores, A., Manosalva, P., Ochoa-Ascencio, S., Díaz-Celaya, M., Rodríguez-Alvarado, G., Fernández-Pavía, S. P., et al. (2022). Characterization and fungicides sensitivity of Phytophthora cinnamomi isolates causing avocado root rot in Zitácuaro, Michoacán. Rev. Mex. Fitopatol. 40, 59– 81. doi: 10.18781/r.mex.fit.2109-4

Mossler, M. A., and Crane, J. H. (2008). Florida Crop/Pest Management Profile: Avocado: CIR 1271/PI048, rev. 9/2008. EDIS 2008. doi: 10.32473/edis-pi048-2008

Ochoa-Fuentes, Y. M., Martínez-de la Vega, O., Olalde-Portugal, V., Cerna-Chávez, E., Landeros-Flores, J., Hernández-Castillo, F. D., et al. (2007). Genetic variability of Phytophthora cinnamomi Rands in Michoacan, Mexico. Rev. Mex. Fitopatol. 25, 161–166.

Ochoa-Fuentes, Y.M., Cerna-Chavez, E., Gallegos-Morales, G., Cepeda-Siller, M., Landeros-Flores, J., and Flores Olivas, A. (2015). Pathogenic variability of Phytophthora cinnamomi Rands in Persea americana Mill. of Michoacan, Mexico. Ecosis. Recur. Agropec. 2, 211–215.

Pagliaccia, D., Pond, E., McKee, B., and Douhan, G. W. (2013). Population genetic structure of Phytophthora cinnamomi associated with avocado in California and the discovery of a potentially recent introduction of a new clonal lineage. Phytopathology 103, 91–97. doi: 10.1094/PHYTO-01-12-0016-R

Palacios-Joya, L. P., Rodríguez-Arévalo, K. A., Martínez, M. F., Murcia-Riaño, N., Rodríguez-Mora, D. M., Palacios-Joya, L. P., et al. (2025). Phytophthora species causing root rot in avocado seedlings at Colombian nurseries: morphological, molecular, and pathogenic analysis. Sci. Agropecu. 16, 113– 121. doi: 10.17268/sci.agropecu.2025.010

Ploetz, R. C., and Parrado, J. L. (1987). Recovery of Phytophthora cinnamomi from avocado soils of South Florida. Proc. Fla. State Hort. Soc. 100, 288–290.

Ploetz, R., and Schaffer, B. (1989). Effects of flooding and Phytophthora root rot on net gas exchange and growth of avocado. Phytopathology 79, 204–208. doi: 10.1094/Phyto-79-204

Ploetz, R. C., Haynes, J. and Schnell, R. J. (2001) Phytophthora root rot-resistant avocado rootstocks for Southern Florida: selection of open-pollinated seedling progeny. Proc. Fla. State Hort. Soc., 114, pp. 6–10.

Qu, T., Shao, Y., Csinos, A. S., and Ji, P. (2016). Sensitivity of Phytophthora nicotianae from tobacco to fluopicolide, mandipropamid, and oxathiapiprolin. Plant Dis. 100, 2119–2125. doi: 10.1094/PDIS-04-16-0429-RE

Ramírez-Gil, J. G., Castañeda-Sánchez, D. A., and Morales-Osorio, J. G. (2017). Production of avocado trees infected with Phytophthora cinnamomi under different management regimes. Plant Pathol. 66, 623–632. doi: 10.1111/ppa.12620

Ramírez-Gil, J. G., and Morales-Osorio, J. G. (2020). Integrated proposal for management of root rot caused by Phytophthora cinnamomi in avocado cv. Hass crops. Crop Prot. 137, 105271. doi: 10.1016/j.cropro.2020.105271

Ramírez-Gil, J. G., Morales-Osorio, J. G., and Peterson, A. T. (2021). The distribution of Phytophthora cinnamomi in the Americas is related to its main host (Persea americana), but with high potential for expansion. Phytopathol. Mediterr. 60, 521–534. doi: 10.36253/phyto-12327

Riley, N. M., Förster, H., and Adaskaveg, J. E. (2024). Regional comparisons of sensitivities of Phytophthora citrophthora and P. syringae causing citrus brown rot in California to four new and two older fungicides. Plant Dis. 108, 1582–1590. doi: 10.1094/PDIS-08-23-1556-RE

Robin, C., and Desprez-Loustau, M.-L. (1998). Testing variability in pathogenicity of Phytophthora cinnamomi. Eur. J. Plant Pathol. 104, 465–475. doi: 10.1023/A:1008649806620

RoyChowdhury, M., Saini, M., Driver, T., and Minton, B. (2016). Distribution of Phytophthora spp. in Southern Texas citrus grove soils. Plant Health Prog. 17, 182–183. doi: 10.1094/PHP-BR-16-0028

Sanchez, A. D., Sosa, M. C., Lutz, M. C., Carreño, G. A., Ousset, M. J., and Lucero, G. S. (2019). Identification and pathogenicity of Phytophthora species in pear commercial orchards in Argentina. Eur. J. Plant Pathol. 154, 811–822. doi: 10.1007/s10658-019-01705-2

Sánchez-González, E. I., Gutiérrez-Soto, J. G., Olivares-Sáenz, E., Gutiérrez-Díez, A., Barrientos-Priego, A. F., and Ochoa-Ascencio, S. (2019). Screening progenies of Mexican race avocado genotypes for resistance to Phytophthora cinnamomi Rands. HortScience 54(5), 809–813. doi: 10.21273/HORTSCI13552-18

Scagel, C. F., Weiland, J. E., Beck, B. R., and Mitchell, J. N. (2023). Temperature and fungicide sensitivity in three prevalent Phytophthora species causing Phytophthora root rot in rhododendron. Plant Dis. 107, 3014–3025. doi: 10.1094/PDIS-11-22-2670-RE

Serrano, M. S., and Garbelotto, M. (2020). Differential response of four Californian native plants to worldwide Phytophthora cinnamomi genotypes: implications for the modeling of disease spread in California. Eur. J. Plant Pathol. 156, 851–866. doi: 10.1007/s10658-020-01936-8

Shakya, S. K., Grünwald, N. J., Fieland, V. J., Knaus, B. J., Weiland, J. E., Maia, C., et al. (2021). Phylogeography of the wide-host range panglobal plant pathogen Phytophthora cinnamomi. Mol. Ecol. 30, 5164–5178. doi: 10.1111/mec.16109

Shands, A. C., Xu, G., Belisle, R. J., Seifbarghi, S., Jackson, N., Bombarely, A., et al. (2024). Genomic and transcriptomic analyses of Phytophthora cinnamomi reveal complex genome architecture, expansion of pathogenicity factors, and host-dependent gene expression profiles. Front. Microbiol. 15. doi: 10.3389/fmicb.2024.1341803

Shands, A. C., Mondragón-Flores, A., Valencia, V., Hua, V., Cauldron, N. C., Grünwald, N. J., et al. (2025). Phytophthora cinnamomi populations affecting avocado in California show low differentiation, phenotypic variability, and introductions from Mexico. MPMI 38, 708–721. doi: 10.1094/MPMI-08-24-0101-R

Swiecki, T. J., Bernhardt, E. A., and McClanahan, S. G. (2024). Validating and optimizing a method for detecting Phytophthora species by baiting leachate from arrays of container nursery plants. PhytoFront. 4, 14–30. doi: 10.1094/PHYTOFR-03-23-0044-FI

Vannini, A., and Morales-Rodriguez, C. (2022). “Phytophthora diseases” in Forest Microbiology, eds. F. O. Asiegbu and A. Kovalchuk (Academic Press), 379–402. doi: 10.1016/B978-0-323-85042-1.00016-1.

Vegh, I., Leroux, P., LeBerre, A. and Lanen, C. (1985). Détection sur Chamaecyparis lawsoniana ‘Elwoodii’ d’une souche de Phytophthora cinnamomi Rands résistante au phoséthyl-Al. PHM Revue Horticole, 262, pp. 19–21.

Vitale, S., Scotton, M., Vettraino, A. M., Vannini, A., Haegi, A., Luongo, L., et al. (2018). Characterization of Phytophthora cinnamomi from common walnut in Southern Europe environment. For. Pathol. 49, e12477. doi: 10.1111/efp.12477

Wager, V. A. (1942). Phytophthora cinnamomi and wet soil in relation to the dying-back of avocado trees. Hilgardia, 14(9), 519–532.

Wilkins, N., Whiley, H., and Ross, K. (2025). A review of the current methods used to detect Phytophthora cinnamomi. Fungal Biol. Rev. 53, 100441. doi: 10.1016/j.fbr.2025.100441

Wilkinson, C. J., Shearer, B. L., Jackson, T. J., and Hardy, G. E. S. J. (2001). Variation in sensitivity of Western Australian isolates of Phytophthora cinnamomi to phosphite in vitro. Plant Pathol. 50, 83–89. doi: 10.1046/j.1365-3059.2001.00539.x

Zentmyer, G. A. (1951). “Avocado diseases in Mexico and Costa Rica,” in California Avocado Society Yearbook 36, 103–104.

Zentmyer, G. A. (1976). “Soil-borne pathogens of avocado,” in Proceedings of the First International Tropical Fruit Short Course: The Avocado, eds. J. W. Sauls, R. L. Phillips, and L. K. Jackson (Gainesville, FL: University of Florida), 75–82.

